# Rhizo-PET: A Dedicated PET System for 4D Imaging of Carbon Dynamics in the Rhizosphere

**DOI:** 10.64898/2026.04.14.718567

**Authors:** Muhammad Nasir Ullah, Daniel Hastings, Seung Joon Lee, Wonu Park, Sarah Jin Zou, David Anders, Jun Hyung Park, Drew Weisenberger, Weixin Cheng, Shiva Abbaszadeh, Craig Levin

## Abstract

Imaging carbon movements in the rhizosphere is fundamentally limited by high soil heterogeneity, low signal levels, and lack of methodology. We present Rhizo-PET, a dedicated positron emission tomography (PET) imaging and analysis framework designed to characterize the 4D spatiotemporal patterns of tracer distribution in intact plant–soil systems. The system achieved a global energy resolution of 11.93 ± 0.02% FWHM at 511 keV and maintained stable performance over 8 h of continuous acquisition, with a coincidence rate variation of only 0.7%. Spatial resolution reached 1.06 mm near the center of the field of view, establishing a high-fidelity region for root-scale analysis. Dynamic datasets were acquired from live *Phaseolus vulgaris* plants (*N*= 3) over 180 min following ^11^CO_2_ pulse labeling and reconstructed into 3 min temporal frames.

Quantitative analysis across 243 independent regions of interest (ROI) revealed that cumulative tracer accumulation decreases monotonically with radial distance from the root axis, while axial transport delays increase systematically in lower root segments (*p <*0.001). Hierarchical variability analysis showed that within-plant spatial organization (*CV*_*TTP*_ = 0.03) is significantly more stable than inter-plant variation (*CV*_*TTP*_ = 0.14), proving that the observed heterogeneity reflects biological spatial organization rather than experimental instability. These results establish Rhizo-PET as a robust, reproducible platform for the non-invasive, time-resolved analysis of carbon dynamics in the rhizosphere under realistic soil conditions.

## 1 Introduction

The rhizosphere—the interface between plant roots and soil—is a major driver of terrestrial carbon cycling, mediating a substantial fraction of global carbon fluxes through root exudation, microbial processing, and mineral stabilization [1–4]. Rhizosphere dynamics are integral to plant physiology, nutrient cycling, and microbial communities from managed croplands to wilderness areas [5]. Rhizosphere processes can account for approximately one third of total carbon and nitrogen mineralization on land [1], despite relatively the small volume of soil—typically defined as 0-2 mm from the root surface [6]. Plants exert strong controls on the rhizosphere through root exudates, which consist of 11–40% of photosynthetically fixed carbon, shape microbial communities, mobilize nutrients and drive the formation of soil aggregates [4, 7, 8].

Photo-assimilated carbon is typically transported in the form of sucrose from the chloroplasts throughout the plant body via the phloem [9]. Less than an hour after photo-assimilation, sucrose may arrive at root tissues [10, 11], where it can be further allocated to various components including: (1) root tissue respiration and active transport; (2) growth and structural support of new root tissues; (3) support of symbionts (e.g., mycorrhizas or rhizobium nodules); (4) bolstering root carbon storage reserves; and (5) biosynthesis of organic compounds including root exudates. Based on the current understanding, transport of exudates across the rhizoplane and into the rhizosphere occurs in different ways depending on the molecular weight of the compound. Low-molecular-weight exudates (e.g. sugars, amino acids) are also classed as “diffusates” as they typically diffuse passively into the soil with the aid of sugar-transport proteins [12]. Sixty to ninety percent of root exudates are low-molecular-weight reducing sugars [11]. Root hairs also use vesicles to passively secrete high-molecular-weight exudates into the rhizosphere [13]. Once exudates have passed the rhizoplane into the rhizosphere, their continued movement and distribution is primarily mediated by diffusion along concentration gradients, advective transport via soil water movement, rapid microbial consumption, and soil water movement [12].

The rhizosphere of each root will has its own set of physical, biological, and chemical conditions giving rise to a tremendous diversity of rhizosphere systems, each with a high level of complexity [14]. Furthermore, these conditions will vary nearly constantly around the root, creating high spatial and temporal heterogeneity even within the same plant. As a result, the rhizosphere is particularly difficult to study, and many models use integrative methods to approximate the overall effect of rhizosphere processes throughout the root [15]. While these approaches have produced important findings, key insights could be obtained by directly observing the spatiotemporal distribution of labeled ^11^C-derived carbon compounds within the rhizosphere.

Methods that can quantitatively measure the *in situ* three-dimensional (3D) distribution and dynamics of labeled ^11^C-derived carbon within this heterogeneous environment over time would be a great boon to rhizosphere science; however, there are many obstacles still to overcome. Direct visualization of photosynthate movement requires imaging approaches that capture both structural and functional information at relevant spatial (*<*2 mm) and temporal (minutes to hours) scales in intact roots and soil systems[16–18]. The soil environment presents unique challenges: its opacity precludes optical methods, its density and heterogeneity complicate X-ray computed tomography (CT), and its three-dimensional (3D) pore architecture hosts processes occurring across multiple scales [19–22]. Pulse-labeling plants with positron-emitting carbon isotopes (^11^C)—a brief exposure to radiolabeled CO_2_—followed by positron emission tomography (PET) enables quantitative and non-invasive tracking of carbon transport. Previous plant-imaging systems such as PETIS, PlanTIS, and early MRI–PET platforms provided pioneering insights into carbon translocation but were fundamentally constrained by fixed planar or partial-ring detector geometries, limited angular sampling, and coarse spatial resolution, yielding only two-dimensional (2D) projections or isolated organ-level snapshots of rhizosphere processes [18, 23–26]. The ability to visualize in four dimensions (4D) the spatiotemporal dynamics of labeled carbon within the rhizosphere represents a major step forward.

To overcome these limitations, we developed *Rhizo-PET (R-PET)*, a dedicated positron emission tomography (PET) platform engineered for 4D live rhizosphere imaging under natural soil conditions (Fig. 1). R-PET features a reconfigurable octagonal detector geometry with active object rotation, achieving complete 3D angular sampling and millimeter spatial resolution across the root–soil interface [27, 28]. This configuration enables real-time, fully 3D, non-invasive visualization of root–soil carbon signal dynamics in intact soil volumes, capturing the spatial and temporal evolution of carbon-associated PET signal patterns.

**Fig. (1).**
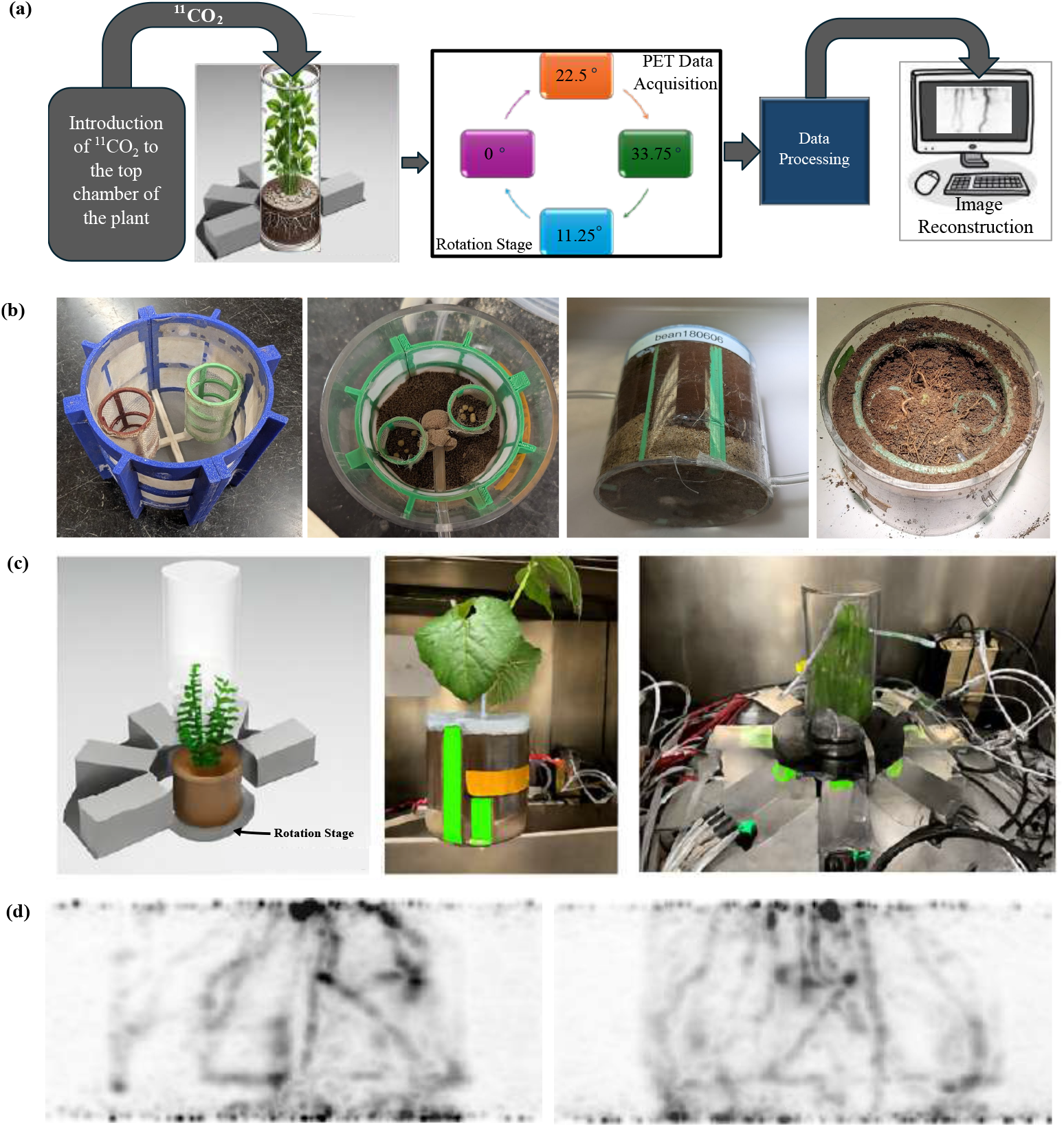
Experimental setup and imaging workflow for Rhizo-PET studies of carbon allocation in plants. (**a**) Schematic of the ^11^C-CO_2_ pulse-labeling and imaging workflow. Radiolabeled CO_2_ was introduced into the top chamber of a live *Phaseolus vulgaris* plant, followed by PET data acquisition at sequential rotation angles (0°, 11.25°, 22.5°, and 33.75°), data processing, and image reconstruction. (**b**) Pot infrastructure, including a plastic frame with 50 micron mesh to prevent roots from being potbound and providing a layer of soil on the opposite side for the flow of water and solutes, upper air flow tube in a soil filled pot, and side and to angles of pot with above-ground portion of the plant removed. The small cylinders within the pot are sub-sampling containers used for a concurrent *µ*CT study (data not shown). (**c**) Photographs and 3D model of the plant holder and custom rotation stage integrated within the PET scanner, showing the assembled setup used for dynamic ^11^C-CO_2_ imaging of the lower chamber. (**d**) Representative reconstructed PET images of ^11^C-CO_2_ movement in the plant–soil system, showing real-time visualization of carbon translocation from shoots to roots and into the rhizosphere.

Using ^11^CO_2_ pulse-labeling, R-PET enables non-invasive tracking of photo-assimilated carbon transport from leaves to roots within intact soil volumes, overcoming the opacity and structural heterogeneity that limit optical, MRI, and X-ray-based approaches. However, quantitative interpretation of rhizosphere PET data presents unique challenges due to spatially varying attenuation, heterogeneous tracer delivery, partial-volume effects at the root–soil interface, and substantial inter-plant variability, which complicate conventional kinetic modeling and absolute activity quantification.

To address these limitations, we characterize R-PET system performance and introduce a non-parametric 4D (x, y, z, t) region-of-interest (ROI) analysis framework designed to extract stable spatiotemporal descriptors from rhizosphere PET data. Rather than relying on absolute tracer uptake values, this framework operates on relative time–activity trends, enabling reproducible comparison of PET signal dynamics associated with labeled ^11^C across roots and surrounding soil regions at 3-minute temporal resolution and millimeter-scale spatial sampling. This approach establishes a robust analytical foundation for quantitative 4D PET studies of carbon dynamics in the rhizosphere.

## 2 Results

### 2.1 Detector performance and acquisition stability

#### 2.1.1 Energy resolution

Following per-crystal photopeak alignment, the system achieved a global energy resolution of 11.93 ± 0.02% FWHM at 511 keV, compared with 15.36 ± 0.28% prior to alignment. Across all scintillator elements, post-alignment energy resolution ranged from 11.93% to 14.11% FWHM, with a mean of 12.24 ± 0.67%, indicating uniform response across detector modules (Fig. 2).

**Fig. (2).**
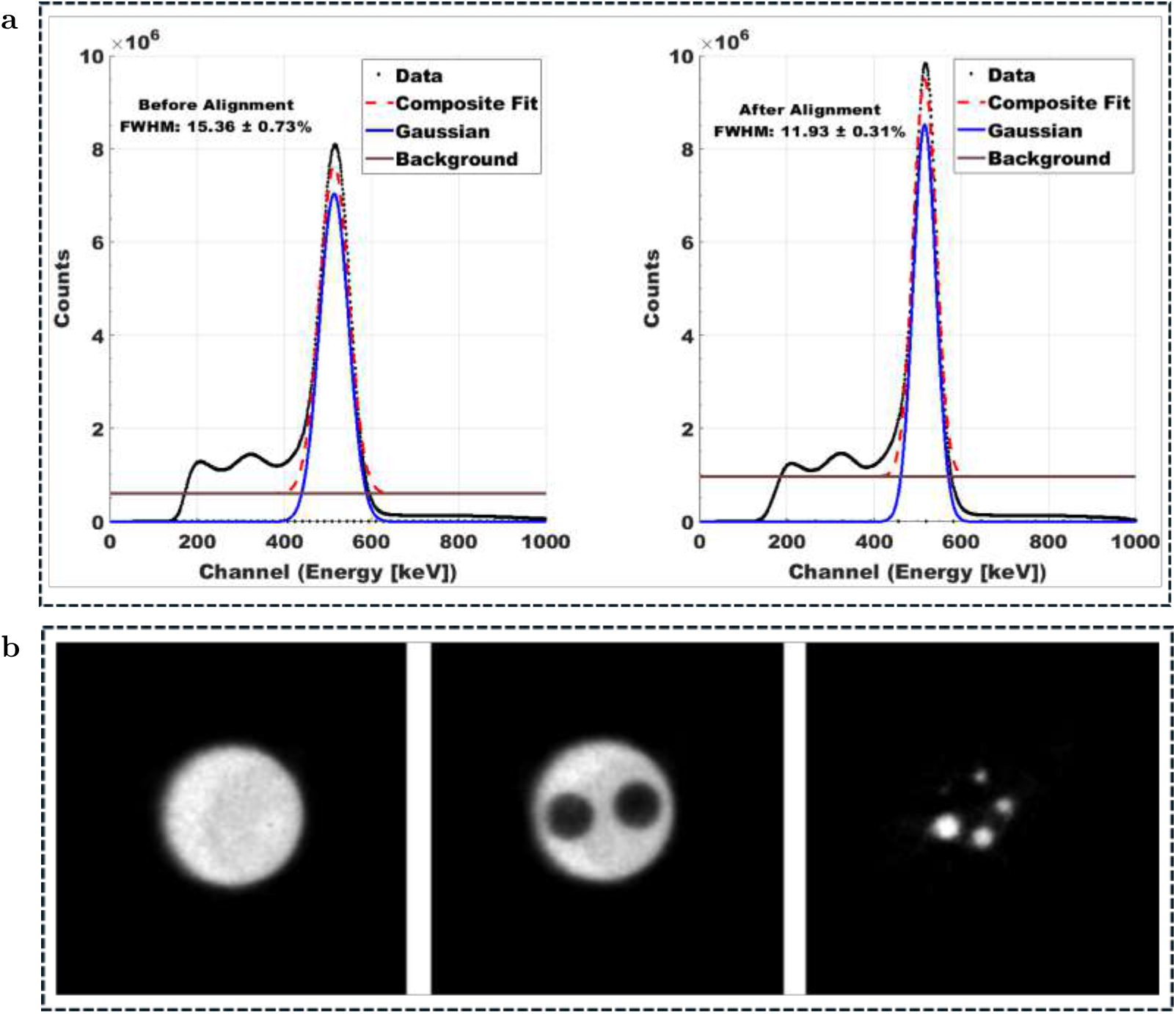
R-PET System Performance Characterization. **(a)** The global energy spectrum of a ^22^Na point source is shown before (left) and after (right) detector-specific photopeak alignment. This alignment improves the global energy resolution from 15.36% to 11.93% FWHM at 511 keV, enabling improved signal quantification via energy-based scatter filtering. **(b)** Reconstruction of the image quality phantom using 3D-MLEM. From left to right, the image shows the uniform region, the air and water chambers, and the hot rod compartment containing 1 mm to 5 mm diameter rods. This image demonstrates the system’s ability to resolve variations in density and feature size.

#### 2.1.2 Temporal stability

Detector stability was evaluated during an 8 h continuous acquisition. Global energy resolution varied by less than ± 0.05% relative to its mean value. The coincidence count rate remained stable at 8.69 × 10^6^ counts min^−1^ with a total variation of 0.7% across 15 min intervals. The mean photopeak shift was − 1.02 keV, indicating minimal gain drift over the acquisition period.

### 2.2 Spatial resolution and image quality

Spatial resolution was assessed using a ^22^Na point source positioned at multiple radial and axial offsets. Tangential and coronal FWHM values remained within approximately 1–2.5 mm near the center of the field of view, while radial resolution degraded with increasing offset, exceeding 13 mm FWHM at 25 mm radial displacement (Table 1). These measurements define the effective region used for subsequent spatial analyses.

**Table (1).** Tangential (T), radial (R), and coronal (C) FWHM and FWTM values. Resolution metrics (in mm) for various axial and radial positions obtained using a ^22^Na point source (250*µ*m active area diameter) and reconstructed with 3D Filtered Back Projection (3DFBP).

| Axial (mm) | Radius (mm) | Tangential (T) |  | Radial (R) |  | Coronal (C) |  |
| --- | --- | --- | --- | --- | --- | --- | --- |
|  |  | FWHM | FWTM | FWHM | FWTM | FWHM | FWTM |
| 0 | 0 | 1.06 | 2.47 | 0.80 | 2.47 | 2.03 | 3.64 |
| 0 | 5 | 1.09 | 2.40 | 0.91 | 2.68 | 2.11 | 3.61 |
| 0 | 10 | 1.14 | 2.01 | 0.80 | 2.63 | 2.07 | 3.57 |
| 0 | 15 | 1.14 | 2.17 | 1.42 | 4.47 | 2.26 | 4.50 |
| 0 | 20 | 1.34 | 2.52 | 13.05 | 15.70 | 2.44 | 5.32 |
| 0 | 25 | 1.34 | 2.40 | 13.65 | 16.76 | 2.28 | 4.73 |
| 12 | 0 | 1.27 | 2.71 | 0.84 | 2.62 | 2.05 | 3.64 |
| 12 | 5 | 1.19 | 2.52 | 0.97 | 2.60 | 2.01 | 3.63 |
| 12 | 10 | 1.29 | 2.09 | 0.58 | 2.57 | 2.09 | 3.70 |
| 12 | 15 | 1.09 | 2.11 | 1.46 | 2.82 | 2.12 | 3.89 |
| 12 | 25 | 1.06 | 2.10 | 14.01 | 25.17 | 2.12 | 4.69 |

**Note:** FWHM = Full Width at Half Maximum; FWTM = Full Width at Tenth Maximum.

Image quality was evaluated using a NEMA NU 4 image-quality phantom. The uniform region exhibited a mean reconstructed signal of 1089.46 ± 54.61 a.u., corresponding to a coefficient of variation of 5.0%. Spillover ratios were 0.097 (air) and 0.158 (water). Recovery coefficients increased monotonically with rod diameter, ranging from 0.201 (1 mm) to 0.869 (5 mm), consistent with measured spatial resolution and partial-volume effects (Table 2).

**Table (2).** Performance metrics for hot rods. Quantitative performance values as a function of rod diameter.

| Rod diameter (mm) | Mean activity | RC | CNR | SBR | Noise (CV%) |
| --- | --- | --- | --- | --- | --- |
| 1 | 218.72 | 0.201 | 1.36 | 13.98 | 68.1 |
| 2 | 490.57 | 0.450 | 1.42 | 31.37 | 68.0 |
| 3 | 618.74 | 0.568 | 1.47 | 39.56 | 66.4 |
| 4 | 806.27 | 0.740 | 1.53 | 51.55 | 63.9 |
| 5 | 946.73 | 0.869 | 1.56 | 60.54 | 62.9 |

**Note:** RC, Recovery Coefficient; CNR, Contrast-to-Noise Ratio; SBR, Signal-to-Background Ratio; CV, Coefficient of Variation.

**Table (3).** Summary of non-parametric kinetic descriptors and hierarchical variability across live plant datasets.

| Descriptor | Intra-root | Inter-root | Inter-plant |
| --- | --- | --- | --- |
| Mean TTP (min) | 74.1 ± 6.2 | 76.8 ± 8.9 | 81.3 ± 12.4 |
| CV(TTP) | 0.03 | 0.06 | 0.14 |
| Mean AUC (a.u. · h) | 1.00 ± 0.24 | 1.00 ± 0.60 | 1.00 ± 0.61 |
| CV(AUC) | 0.24 | 0.60 | 0.61 |
| ROIs analyzed | 81 | 243 | 243 |

### 2.3 Live plant imaging results

Dynamic PET imaging of live plants produced ROI-level time-activity curves across roots and spatial locations which describe the heterogeneity of the rhizosphere [14]. (Fig. 3). Despite large differences in signal amplitude, many ROIs exhibited well-defined temporal structure, enabling consistent extraction of timing- and accumulation-based descriptors.Detailed ROI-level metrics across all independent plant datasets are provided in the supplementary materials (Table. S2).

**Fig. (3).**
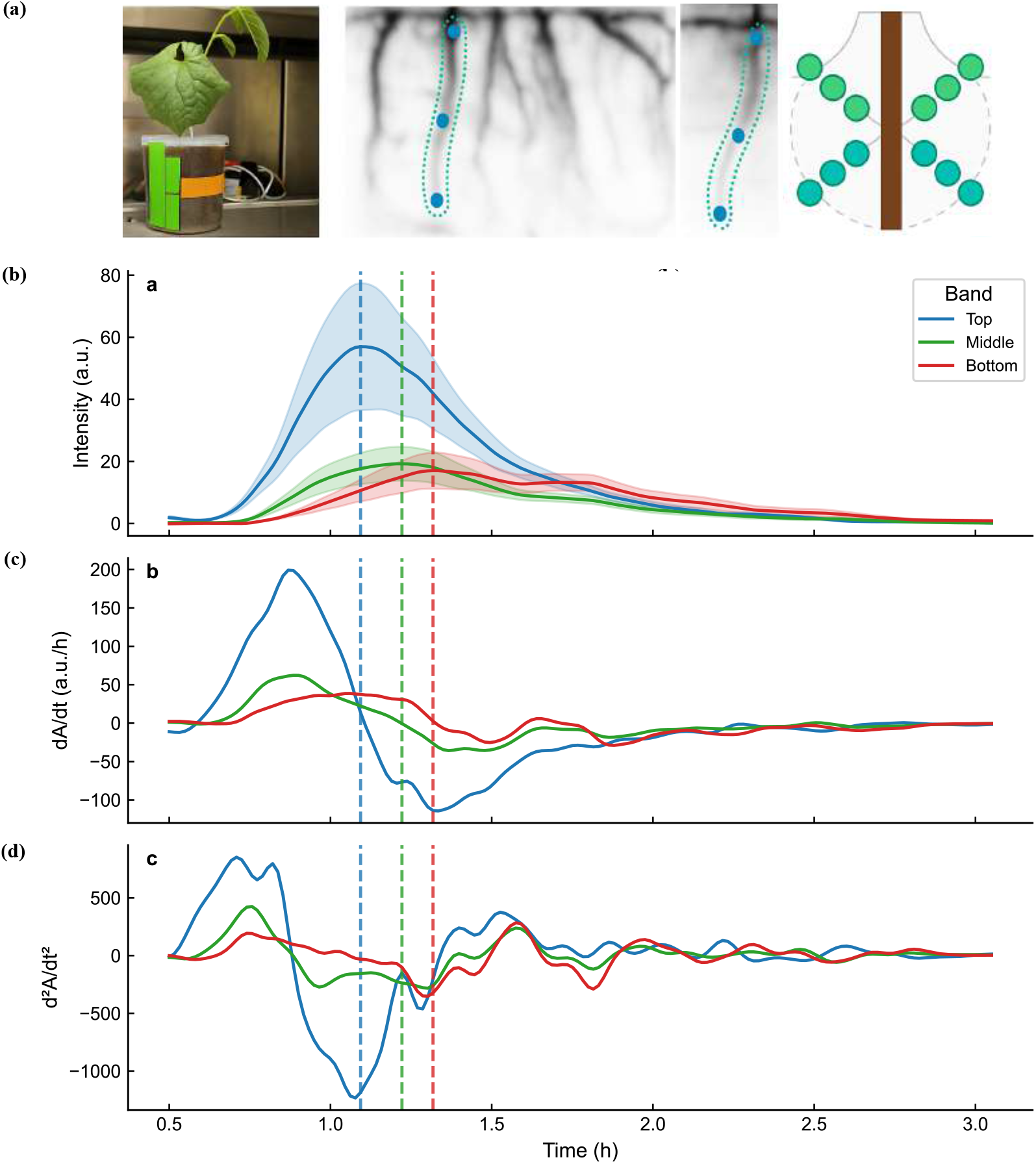
Experimental setup and band-resolved temporal analysis of root PET signal. (a) Photograph of the potted plant during imaging, representative root projection with extracted root centerline and segmented regions of interest (top, middle, bottom), and a schematic illustrating the band definition along the root axis. (b) Mean PET signal intensity (a.u.) versus time for each band (shaded region indicates variability); dashed vertical lines denote the characteristic peak time for each band. (c) First temporal derivative of the activity curves, *dA/dt*, highlighting differences in uptake/transport dynamics along the root. (d) Second temporal derivative, *d*^2^*A/dt*^2^, emphasizing acceleration and inflection points in tracer transport.

### 2.4 Spatial organization of tracer accumulation

Across all plants, cumulative tracer accumulation decreased monotonically with increasing radial distance from the root axis (Fig. 4a,c). Although absolute AUC (area under the curve) values varied between plants, relative spatial ordering was preserved, indicating reproducible radial organization Table. S2). Variance decomposition showed that variability within plants exceeded inter-plant variability (Fig. 4a), suggesting that observed heterogeneity primarily reflects spatial structure rather than experimental instability.

**Fig. (4).**
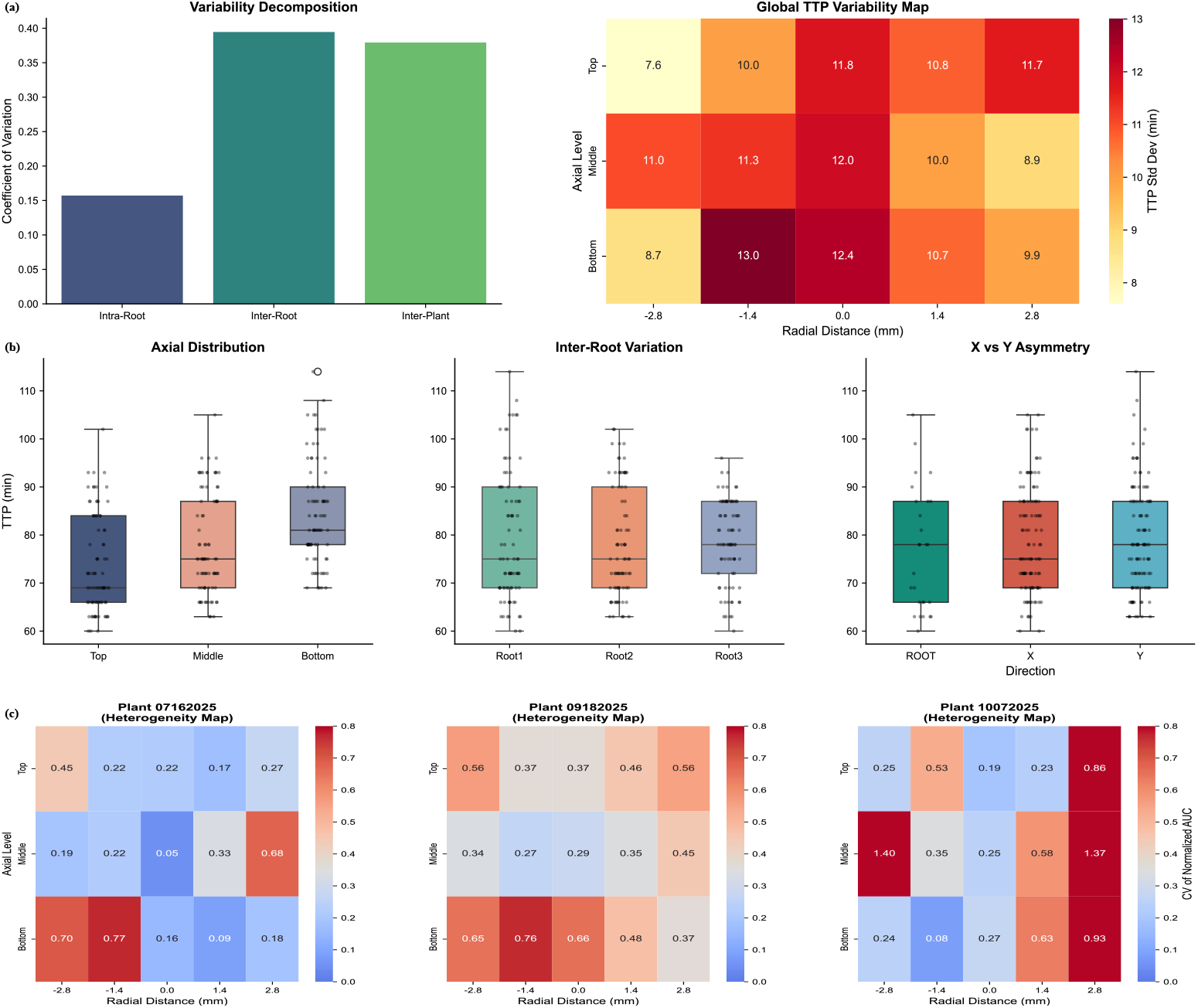
(a) Variability decomposition of non-parametric descriptors across hierarchical scales. Bars show mean coefficient of variation (CV) for time-to-peak (TTP) and area-under-the-curve (AUC) computed across intra-root (orthogonal X vs Y directions), inter-root (within plant), and inter-plant levels, demonstrating increasing variability with hierarchical scale. (b) Spatial and directional structure of TTP across all plants and roots. Left: axial TTP distributions (Top, Middle, Bottom) pooled across datasets. Center: inter-root variability within individual plants. Right: comparison of TTP distributions along orthogonal X and Y directions, indicating minimal directional bias. (c) Per-plant spatial heterogeneity maps of cumulative tracer accumulation. Heatmaps show the coefficient of variation (CV) of decay-corrected AUC values for each plant as a function of radial distance from the root axis and axial position (Top, Middle, Bottom), revealing reproducible spatial heterogeneity patterns across independent datasets.

**Fig. (5).**
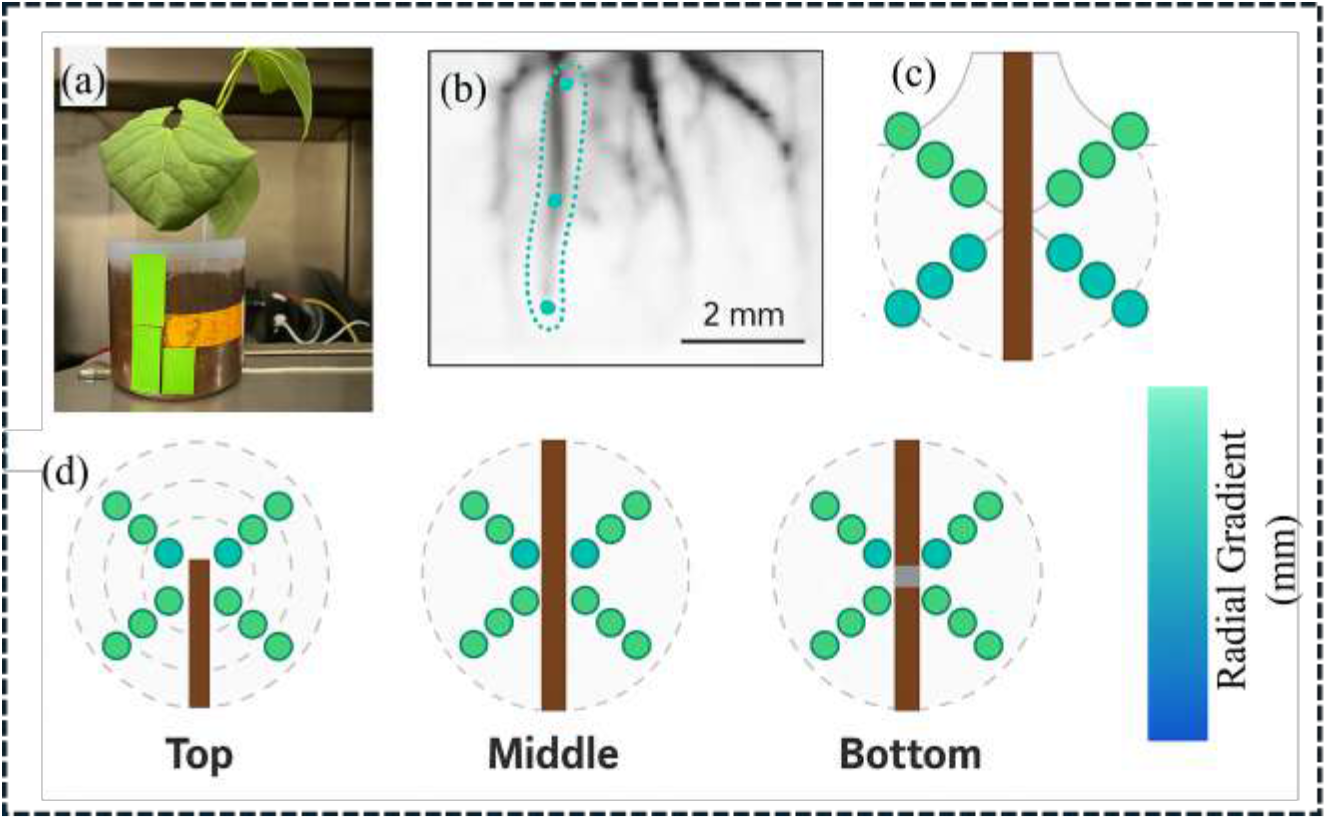
Definition of axial and lateral regions-of-interest (ROIs) for Rhizo-PET kinetic analysis. **(a)** Photograph of a live *Phaseolus vulgaris* plant during ^11^CO_2_ pulse labeling and PET acquisition. **(b–c)** Reconstructed PET images illustrating axial ROIs along the primary root at multiple depths. **(d)** Schematic representation of the corresponding ROI placement along the main root axis. **(e–g)** Cross-sectional schematics depict lateral ROIs extending from the root surface into surrounding soil at the top, middle, and bottom sections. **(h)** Lateral gradient scale (mm) consistently applied across kinetic analyses. This ROI framework enabled extraction of time–activity curves (TACs), time-to-peak (TTP), and area-under-the-curve (AUC) metrics for microscale quantification of carbon allocation dynamics in the rhizosphere.

### 2.5 Transport timing and spatial gradients

Temporal transport metrics revealed systematic delays along the axial direction, with later arrival times observed in lower root segments (Fig. 3b). This axial delay was statistically significant (*p <* 0.001, Table. S3). Radially, the TTP (Time-to-Peak) values were shortest near the root center and increased toward the periphery, forming a shallow but consistent radial gradient (Fig. 4b); Table. S3). No significant directional asymmetry was observed between orthogonal sampling directions (*p* = 0.86 Table S3) and (Fig. 4b).

### 2.6 Hierarchical variability

Temporal variability increased with hierarchical scale, progressing from intra-root to inter-root and inter-plant comparisons (Fig. 4a). Spatial maps of timing and accumulation variability revealed elevated dispersion at intermediate radial distances and in lower axial segments (Fig. 4c), patterns that were conserved across independent plant datasets.

### 2.7 Robustness and reproducibility

Spatial ordering of ROIs was preserved across independent datasets and varying signal conditions, as demonstrated by the consistent axial and radial metrics (Table. S2). The high reproducibility of these spatial patterns across different roots and plants (Table. S4), combined with their high statistical significance (*p* < 0.001, Table. S3), confirms that the observed patterns arise from coherent transport dynamics rather than noise or smoothing artifacts.

## 3 Materials and Methods

### 3.1 Rhizo-PET (R-PET) system architecture

Rhizo-PET (R-PET) is a custom-built positron emission tomography (PET) system used to acquire time-resolved coincidence data from plant root systems contained within soil-filled vessels. The system consists of eight detector modules arranged in a fixed octagonal geometry, providing an active radial field-of-view (FOV) of 160 mm and an axial FOV of 48 mm (Fig. 1c; Fig. S2). The detector geometry remains fixed throughout data acquisition. Angular sampling is increased exclusively through rotation of the imaging object.

Each detector module incorporates a Hamamatsu H8500 flat-panel position-sensitive photomultiplier tube (PSPMT) with a 49 mm × 49 mm active photocathode and 64 anodes arranged in an 8 × 8 matrix [28]. The PSPMT external dimensions are 52 mm × 52 mm × 27.4 mm. Signals from the 64 anodes are combined using a four-channel resistive charge-division network and amplified using custom front-end electronics prior to digitization [29].

Each PSPMT is optically coupled to a LYSO:Ce scintillator array (Proteus Inc.) consisting of 48 × 48 discrete crystal elements. Individual crystals measure 1.0 mm × 1.0 mm × 10 mm and are arranged with a 1.0 mm pitch. No depth-of-interaction encoding is implemented.

### 3.2 Data acquisition electronics

Signals from the eight detector modules are digitized using two EFADC-16 Ethernet-based data acquisition units developed at Jefferson Lab [29]. Each EFADC-16 unit digitizes 16 input channels at 256 MSPS using 12-bit flash analog-to-digital converters (ADCs). On-board field-programmable gate arrays perform real-time threshold discrimination and coincidence sorting.

The two EFADC-16 units are synchronized via optical fiber to maintain temporal alignment across all channels. Event data are streamed to a host computer over 1000 Base-T Ethernet and stored in list-mode format. Only prompt coincidence events are recorded; no delayed coincidence window is employed.

### 3.3 Acquisition parameters

All PET data were acquired in non–time-of-flight (non-TOF) coincidence mode. A fixed coincidence timing window of 12 ns was applied for all measurements. Events were accepted only if both detected photons fell within an energy window of 450–600 keV. These acquisition parameters were applied identically for system characterization, normalization, and live plant imaging.

For live plant experiments, list-mode data were acquired continuously for a total duration of 180 min following completion of the ^11^CO_2_ pulse-labeling period. Dynamic image reconstruction was performed exclusively using 3-minute temporal frames. No alternative temporal binning was used at any stage of processing or analysis.

### 3.4 System characterization

#### 3.4.1 Rotational acquisition

To increase angular sampling, the imaging object was mounted on a motorized rotation stage positioned concentrically within the detector ring (Fig. 1c). Data were acquired at four discrete angular positions: 0^°^, 11.25^°^, 22.5^°^, and 33.75^°^ (Fig. 1a). At each position, list-mode data were acquired for 30 s prior to rotation to the next angle. Rotation angles and timestamps were recorded and used for angle-tagged rebinning during reconstruction.

#### 3.4.2 Energy resolution and temporal stability

Energy resolution and temporal stability were evaluated using a ^22^Na point source (250 *µ*m active diameter, 1 MBq). Data were acquired continuously for 8 h. Energy spectra were aligned post-acquisition using per-detector linear gain correction. Each spectrum was modeled using a Gaussian photopeak with a linear background,

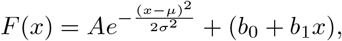

and energy resolution at 511 keV was reported as full width at half maximum (FWHM),

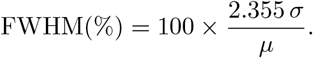

#### 3.4.3 Spatial resolution

Spatial resolution was evaluated using the same ^22^Na point source positioned at radial offsets of 5, 10, 15, and 25 mm and at axial positions *Z* = 0 mm and *Z* = 12 mm. Data were reconstructed using three-dimensional filtered backprojection (3D-FBP) with identical voxelization to experimental data. Radial, tangential, and axial resolutions were quantified using full width at half maximum (FWHM) and full width at tenth maximum (FWTM) following NEMA NU-4–2008 guidelines.

#### 3.4.4 Normalization and reconstruction

Detector sensitivity normalization was performed using a custom annulus phantom filled with 25 mL of a 14.8 MBq ^18^F solution. All data were processed using the same 450–600 keV energy window applied during imaging. Image reconstruction was performed using maximum-likelihood expectation–maximization (MLEM) with 25 iterations and an isotropic voxel size of 0.5 mm × 0.5 mm × 0.5 mm.

No attenuation, scatter, or randoms correction was applied. Reconstructed volumes therefore represent relative coincidence count density rather than absolute activity concentration.

### 3.5 Live plant preparation, PET imaging, and ROI-based analysis

#### 3.5.1 Plant growth and soil preparation

Three independent (*N* = 3) *Phaseolus vulgaris* L. plants were cultivated in cylindrical acrylic pots (100 mm diameter × 100 mm height). Pots were filled with field soil collected from the Ap horizon of a sandy–loam Mollisol from Santa Cruz, California (fine-loamy, mixed, thermic Pachic Agrizerolls). Soil was air-dried, passed through a 2 mm sieve, homogenized, and had a bulk density of 1.25 g cm^−3^. Gravimetric water content at the time of imaging was maintained between 15-20%. To enable soil aeration, two opposing air-flow tubes (1/8 inch inner diameter) were inserted laterally into each pot. Air was circulated through the soil for 15 min every 4 h at approximately 600 mL min^−1^. Root growth was confined to a central cylindrical volume (∼ 320 mL) using a custom 3D-printed PLA insert incorporating a 50 *µ*m mesh, which allowed the flow of solutes, root hairs and mycorhizae and prevented roots from growing against the side of the pot. Plants were grown for 20-25 days under controlled greenhouse conditions (25^°^C day, 18^°^C night).

#### 3.5.2 ^11^CO_2_ pulse labeling and PET acquisition

Above- and below-ground compartments were sealed using food-grade silicone to prevent gas exchange during tracer administration. Plants were illuminated using a 400 W LED source providing approximately 1500 *µ*mol m^−2^ s^−1^ at the canopy. ^11^CO_2_ was produced using a GE PETtrace 880 cyclotron and introduced into a sealed acrylic chamber containing the above-ground portion of the plant. A total activity of 550 MBq *±* 30 MBq was introduced to the chamber and allowed to integrate over a 20 min pulse period. Residual tracer was removed using an alkaline CO_2_ sorbent prior to reopening the chamber to ambient air.

Dynamic PET acquisition of the below-ground compartment began 30 min post-injection and continued for 180 min. Dynamic images were reconstructed into contiguous 3 min temporal frames. All reported times correspond to time post-injection; the initial 30 mins were not used due to high levels of noise from the ^11^CO_2_ introduction. *(Analysis scripts explicitly account for this acquisition offset*.)

#### 3.5.3 Geometry-based ROI definition and TAC extraction

ROIs were defined using a geometry-driven procedure applied identically across all plants and roots (Fig. 3). For each plant, three primary roots were selected based on visual continuity across multiple reconstructed axial slices and exclusion of roots intersecting the pot boundary. Root selection was performed by a single operator following a fixed protocol and was completed prior to TAC extraction; selection was not informed by signal intensity, TAC shape, or downstream metrics.

For each selected root, three axial segments were defined along the visible root length (Top, Middle, Bottom). Within each axial segment, a set of voxel-matched cubic ROIs (0.5 mm × 0.5 mm × 0.5 mm) was placed at fixed radial distances from the root centerline in orthogonal X and Y directions. In the analysis pipeline, radial sampling was performed at − 2.8, − 1.4, 0, +1.4, and +2.8 mm from the root center in each direction, yielding 30 ROIs per root (3 axial levels × 2 directions × 5 radii) and 90 ROIs per plant.

Each ROI was assigned a unique label encoding axial level, direction, and radial distance (e.g., `Top_X_+1.4mm`). TACs were extracted by averaging voxel intensities within each ROI for each 3 min frame.

#### 3.5.4 TAC preprocessing, kinetic descriptors, and variability metrics

All downstream analysis was performed in Python (NumPy, SciPy, pandas, matplotlib). TACs were smoothed using Gaussian filtering (*σ* = 2 time points), and first and second temporal derivatives were computed by numerical differentiation to derive uptake and acceleration profiles.

##### Decay correction

For intensity-based comparisons (peak intensity and AUC), TACs were corrected for ^11^C radioactive decay (*t*_1*/*2_ = 20.334 min) as

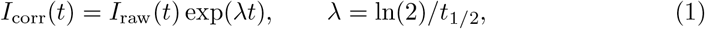

where *t* is time post-injection (not time since imaging start).

##### Timing metrics from non-decay-corrected kinetics

TTP and other timing descriptors were computed from the smoothed *non-decay-corrected* TACs to avoid exponential amplification of late frames that can introduce artificial late maxima (for ^11^C, the correction factor becomes very large by 180 min). Operationally, derivatives (and TTP) were computed from the smoothed intensity series, while AUC was computed from the decay-corrected series.

From each ROI TAC, we extracted: (i) TTP (time of maximum smoothed intensity), (ii) peak intensity, (iii) maximum uptake rate and its time (peak of first derivative), (iv) inflection time (zero-crossing of second derivative), (v) washout rate (mean negative slope after peak), and (vi) AUC (trapezoidal integral of the decay-corrected TAC).

##### Hierarchical heterogeneity quantification

Biological heterogeneity was summarized at three spatial/biological scales: intra-root (directional asymmetry, X vs Y at matched radii), inter-root (variation among roots within a plant), and inter-plant (variation among plants). For mass-accumulation heterogeneity maps, AUC values were normalized within each root (AUC at each ROI divided by total root AUC) prior to computing coefficients of variation (CV) across roots at matched axial/radial positions.

#### 3.5.5 Statistical testing and uncertainty estimation

Differences in TTP across axial levels (Top/Middle/Bottom) were tested using a Kruskal–Wallis H-test. Directional asymmetry (X vs Y) was tested using a Mann– Whitney U test. Radial gradients in timing were assessed using linear regression. Variance components were estimated to partition total TTP variance across hierarchical levels, and bootstrap confidence intervals (95%, *n* = 1000 resamples) were computed for mean estimates.

## 4 Discussion

This study establishes Rhizo-PET as an integrated experimental and analytical framework for resolving relative spatiotemporal patterns of tracer distribution in intact plant–soil systems. The system documented the movement of photosynthate through the roots and into the soil as exudates and captured the variation in this movement in high spatial and temporal resolution. This approach emphasizes reproducible temporal structure and spatial ordering of PET-derived signals rather than pursuing absolute activity quantification, reflecting the heterogeneous attenuation and scattering conditions inherent to soil environments [18, 30, 31].

System characterization demonstrated stable detector performance over multi-hour acquisitions. Energy resolution, coincidence-rate stability, and temporal consistency (Fig. 2, Tables 1–2) support reliable extraction of region-level time–activity curves (TACs), providing a robust instrumental basis for analyses focused on relative timing and accumulation descriptors.

Across all plant datasets, spatial organization of tracer accumulation was consistently observed. Cumulative uptake decreased monotonically with increasing radial distance from the root axis, while axial transport delays increased from upper to lower root segments (Fig. 4b,c). This axial delay was statistically validated (*p <*0.001, Table. S3). Although absolute signal levels varied between experiments, relative spatial ordering was preserved, indicating reproducible organization across independent datasets (Table. S2).

Hierarchical variability analysis showed that within-plant variability exceeded inter-plant variability (Fig. 4a; Table 3). Detailed inter-root and inter-plant TTP metrics, including standard deviations across the 0.0 mm to *pm* 2.8 mm radial range, are documented in Tables. S2 and .S4 This result indicates that the dominant source of dispersion arises from complexity in individual plant-soil systems rather than from experimental instability or dataset-specific effects.

The geometry-based ROI protocol, applied identically across datasets and independent of signal intensity, contributes to interpretability and reproducibility. While manual root selection remains a limitation, the fixed protocol and rank-based analysis mitigate subjective bias and provide a transparent baseline for future extensions incorporating automated segmentation or voxelwise analysis.

Several limitations warrant consideration. The absence of random, scatter, and attenuation correction precludes absolute activity estimation and direct comparison with conventional kinetic models. In addition, spatial resolution degrades at large radial offsets, restricting uniform sensitivity to a central field-of-view region. Finally, the present study includes a limited number of plants, and biological interpretation is therefore intentionally conservative.

Overall, these results demonstrate that dynamic PET imaging can yield reproducible, spatially structured information in the rhizosphere when analysis is explicitly aligned with the physical limits of the measurement. Spatial ordering of ROIs was preserved across all independent plant replicates, as evidenced by the consistent axial and radial metrics reported in the methodological validation tables (Table. S2, Table. S4).

## 5 Conclusion

Rhizo-PET integrates a purpose-built PET system with a non-parametric, ROI-based analysis pipeline to enable reproducible, time-resolved characterization of tracer dynamics in intact plant–soil systems. The results demonstrate that relative spatial ordering and temporal structure of PET signals can be consistently extracted and compared across independent datasets without reliance on absolute activity calibration. This framework establishes a rigorous methodological baseline for rhizosphere PET studies and provides a platform for future advances in correction strategies, segmentation, and multimodal integration.

## Acknowledgements

We acknowledge the Cyclotron Research Facility at Stanford University and the UCSC Greenhouse Facilities for providing resources and technical assistance. We also thank Ricky Padmore for his help with the plant growth apparatus and related components, and Derek Innes (MIPS, Stanford University) for his support in mechanical design.

## Declarations

### Ethics approval and consent to participate

N/A

### Consent for publication

N/A

### Data availability

N/A

### Materials availability

N/A

### Code availability

Code is available at….

### Author contribution

Conceptualization: WC, CL, SA, MNU, DOH Software: SJZ, SJL, MNU, DW Resources: SJL, DW Formal analysis: MNU, WP, SJZ Investigation: MNU, DOH Writing – original draft: MNU, DOH Writing – review & editing: WC, CL, SA Visualization: MNU, WP Supervision: SA, WC, CL Project administration: SA Funding acquisition: SA, WC, CL Validation: MNU DOH

