## Supplemental Data for "Rhizo-PET: A Dedicated PET System for 4D Imaging of Carbon Dynamics in the Rhizosphere"

### Supplementary Data

**Table (S1). Spill-over ratio (SOR) in different media.** Values calculated using 3D-MLEM reconstruction of the NEMA NU 4 phantom.

| Medium | Reconstruction method | SOR |
|---|---|---|
| Water | 3D-MLEM | 0.158 |
| Air | 3D-MLEM | 0.097 |

**Note:** SOR, spill-over ratio; three-dimensional maximum likelihood expectation maximization (3D-MLEM).

**Table (S2).** Inter-Plant Variability for TTP: Methodological validation across all axial levels and radial distances.

| Axial Level | Direction | Radial Dist (mm) | Mean TTP ± SD (min) | CV | N (Plants) |
|---|---|---|---|---|---|
| Bottom | ROOT | 0.0 | 83.33 ± 12.66 | 0.15 | 3 |
| Bottom | X | -2.8 | 85.67 ± 8.33 | 0.10 | 3 |
| Bottom | X | -1.4 | 84.67 ± 11.59 | 0.14 | 3 |
| Bottom | X | 1.4 | 84.00 ± 11.00 | 0.13 | 3 |
| Bottom | X | 2.8 | 85.33 ± 9.61 | 0.11 | 3 |
| Bottom | Y | -2.8 | 82.33 ± 8.39 | 0.10 | 3 |
| Bottom | Y | -1.4 | 84.00 ± 13.45 | 0.16 | 3 |
| Bottom | Y | 1.4 | 84.33 ± 12.06 | 0.14 | 3 |
| Bottom | Y | 2.8 | 87.33 ± 8.39 | 0.10 | 3 |
| Middle | ROOT | 0.0 | 75.33 ± 13.32 | 0.18 | 3 |
| Middle | X | -2.8 | 80.33 ± 14.47 | 0.18 | 3 |
| Middle | X | -1.4 | 76.00 ± 13.23 | 0.17 | 3 |
| Middle | X | 1.4 | 77.33 ± 10.21 | 0.13 | 3 |
| Middle | X | 2.8 | 80.33 ± 7.77 | 0.10 | 3 |
| Middle | Y | -2.8 | 81.00 ± 9.54 | 0.12 | 3 |
| Middle | Y | -1.4 | 76.33 ± 12.86 | 0.17 | 3 |
| Middle | Y | 1.4 | 77.33 ± 12.10 | 0.16 | 3 |
| Middle | Y | 2.8 | 78.67 ± 11.24 | 0.14 | 3 |
| Top | ROOT | 0.0 | 71.00 ± 13.00 | 0.18 | 3 |
| Top | X | -2.8 | 74.00 ± 8.89 | 0.12 | 3 |
| Top | X | -1.4 | 72.33 ± 12.86 | 0.18 | 3 |
| Top | X | 1.4 | 74.00 ± 12.12 | 0.16 | 3 |
| Top | X | 2.8 | 74.00 ± 10.00 | 0.14 | 3 |

**Table (S2).** Inter-Plant Variability for TTP (continued)

| Axial Level | Direction | Radial Dist (mm) | Mean TTP ± SD (min) | CV | N (Plants) |
|---|---|---|---|---|---|
| Top | Y | -2.8 | 77.33 ± 7.57 | 0.10 | 3 |
| Top | Y | -1.4 | 73.67 ± 9.07 | 0.12 | 3 |
| Top | Y | 1.4 | 72.33 ± 12.74 | 0.18 | 3 |
| Top | Y | 2.8 | 76.67 ± 14.84 | 0.19 | 3 |

*Note: SD (Standard Deviation) is calculated as $Mean \times CV$ for each grouping.*

**Table (S3).** Summary of Statistical Validation Tests.

| Test Scope | Statistical Method | Test Statistic | P-Value |
|---|---|---|---|
| Axial (Top vs Mid vs Bottom) | Kruskal-Wallis | 37.99 | $5.63 \times 10^{-9}$ |
| Directional (X vs Y) | Mann-Whitney U | 5752.0 | 0.862 |
| Radial (0.0 to 2.8 mm) | Linear Regression | $F = 1.587$ | 0.141 |
| Hierarchy (Between Plant) | Variance Component | 0.949 | – |

*Note: All tests were performed on the raw TTP data extracted from the 243 independent ROIs.*

**Table (S4).** Inter-Root Variability: Summary of TTP consistency within each independent plant.

| Plant ID | Axial Level | Mean TTP ± SD (min) | CV | Pairwise Diff | N Roots |
|---|---|---|---|---|---|
| 07162025 | Bottom | 81.00 ± 5.20 | 0.06 | 6.0 | 3 |
| 07162025 | Middle | 72.00 ± 6.00 | 0.08 | 8.0 | 3 |
| 07162025 | Top | 63.00 ± 3.00 | 0.05 | 2.0 | 3 |
| 09182025 | Bottom | 72.00 ± 5.20 | 0.07 | 6.0 | 3 |
| 09182025 | Middle | 64.00 ± 1.73 | 0.03 | 2.0 | 3 |
| 09182025 | Top | 64.00 ± 1.73 | 0.03 | 2.0 | 3 |
| 10072025 | Bottom | 97.00 ± 9.17 | 0.09 | -12.0 | 3 |
| 10072025 | Middle | 90.00 ± 3.00 | 0.03 | -2.0 | 3 |
| 10072025 | Top | 86.00 ± 6.24 | 0.07 | -2.0 | 3 |

*Note: These metrics validate that axial delays are reproducible within the internal root structure of each plant.*

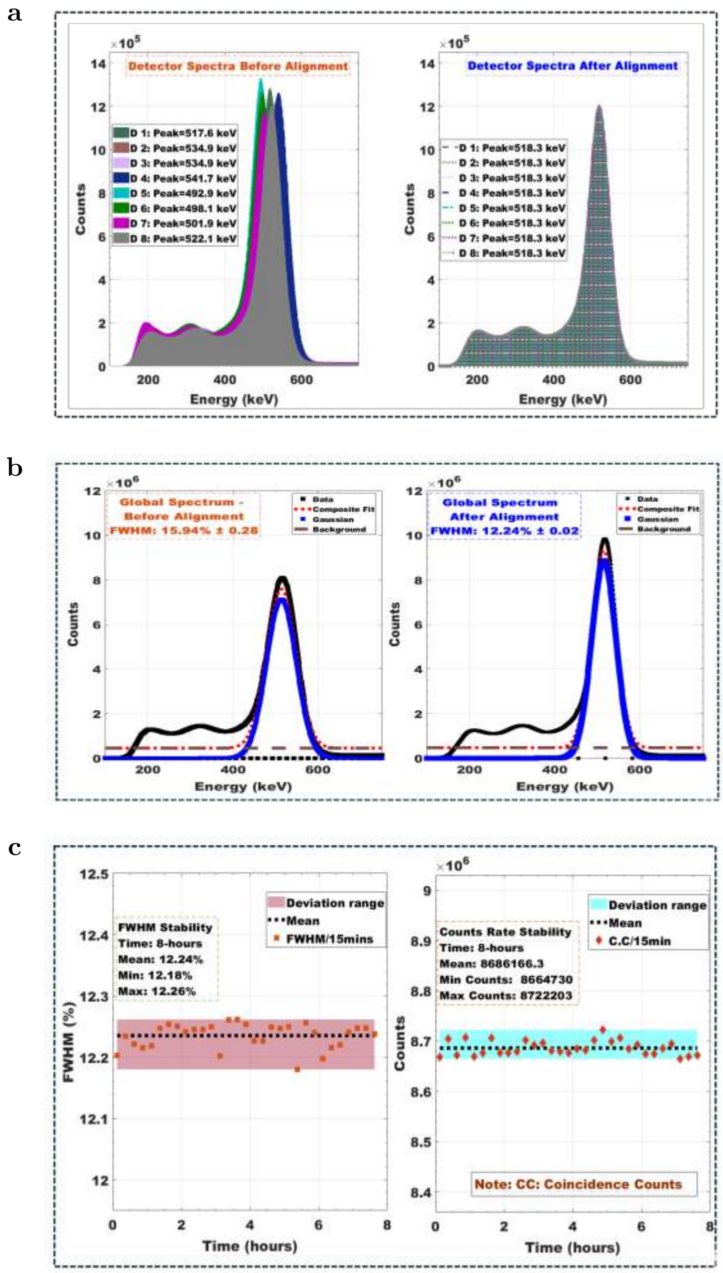


**Fig. (S1). (a) Energy spectra from individual detectors before and (b) after 511 keV photopeak alignment, showing variations due to PSPMT gain differences and improved spectral consistency across detectors.** (c) Comparison of global energy spectra before and (d) after alignment, with measured energy resolution improvements. (e) Energy resolution (FWHM) and (f) coincidence count rate stability over an 8-hour acquisition period.

**Table (S5).** System architecture specifications of the Rhizo-PET (R-PET) scanner used in this study. All values correspond to the fixed configuration employed for data acquisition.

| **Parameter** | **Specification** |
|---|---|
| Scintillator material | LYSO:Ce |
| Scintillator array | 48 × 48 discrete crystal elements |
| Crystal dimensions ($mm^3$) | 1.0 × 1.0 × 10.0 |
| Crystal pitch (mm) | 1.0 |
| Photodetector | Hamamatsu H8500 position-sensitive PMT |
| Active photodetector area ($mm^2$) | 49 × 49 |
| Detector modules | 8 |
| Detector geometry | Fixed octagonal ring |
| Total number of crystals | 18,432 |
| Radial field of view (mm) | 160 |
| Axial field of view (mm) | 48 |
| Depth-of-interaction encoding | None |
| Signal readout | Four-channel resistive charge division |

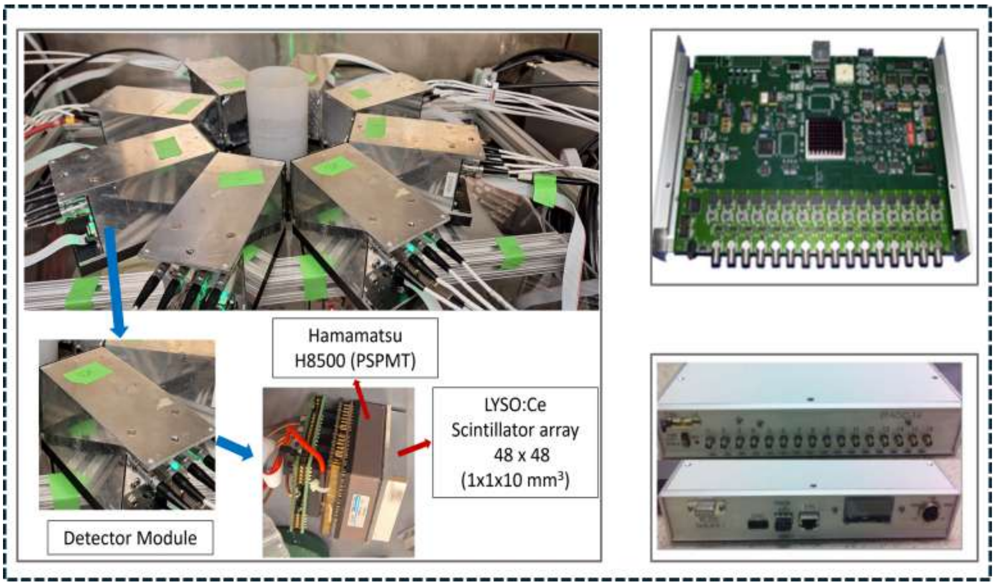


**Fig. (S2). System geometry and detector design of the Rhizo-PET (R-PET) scanner.** The R-PET system comprises eight detector modules arranged in an octagonal geometry (top), optimized for high-sensitivity dynamic imaging of plant roots and rhizosphere processes. The lower left panel shows an individual detector module consisting of a position-sensitive photomultiplier tube (PSPMT, Hamamatsu H8500) optically coupled to a Ce:LYSO scintillator array. Each crystal element measures 1 × 1 × 1 $mm^3$, and the array is matched to the 50 × 50 $mm^2$ active area of the PSPMT. The right panel depicts the custom acquisition board, and the lower right image shows the assembled data acquisition electronics used for synchronized readout of all detector modules.
